# Robust taxonomic classification in gut and vaginal microbiomes demonstrated through benchmarking with age-specific synthetic communities

**DOI:** 10.64898/2026.07.06.736764

**Authors:** Julian M. Trachsel, Hannah Sturgeon, David Goad, Ruben A. Mars, Cheryl Sew Hoy, Kimberley V. Sukhum

## Abstract

Accurate taxonomic profiling of human microbiomes is essential for advancing research and understanding the complex role microbial communities play in human health. When using shotgun metagenomics, the sequencing data is analyzed through metagenomic pipelines, which incorporate various open-source tools and classify microbes based on matched paired-end DNA reads. However, differences in sequencing and computational approaches can produce substantially different microbiome profiles from the same sample, making validation critical. One approach for validation is benchmarking with realistic mock communities, but this remains relatively rare. Additionally, existing benchmarks often overlook microbiome variability across life stages and body sites, limiting their clinical and research utility. Here, we developed age- and body site-stratified synthetic metagenomes, enabling context-aware benchmarking of microbiome pipelines.

Using novelty-based sampling to prioritize microbial diversity and minimize redundancy among selected samples, we selected 300 representative, real biological samples spanning six categories: adult, child, toddler, and infant (>6 months and <6 months) gut samples, as well as adult vaginal samples. We validated three pipelines, Tiny Health’s proprietary Metagenomic Classifier v2 (THMCv2), MetaPhlAn4, and Kraken2+Bracken, using precision, recall, F1 score, and area under the precision-recall curve (AUPR) across age groups and sample types. THMCv2 demonstrated higher recall and F1 scores, detecting more taxa across sample types and ages, while MetaPhlAn4 achieved the highest precision. THMCv2 also achieved the highest area under the precision-recall curve, reflecting peak performance across both abundant and rare species. When analyses were weighted by abundance, THMCv2 and MetaPhlAn4 each characterized the mock community nearly perfectly. Errors for THMCv2 were largely restricted to very low-abundance taxa (<0.001%), whereas MetaPhlAn4 occasionally produced false positives for higher-abundance taxa. Species-level analyses of clinically relevant microbes confirmed these patterns, with THMCv2 demonstrating higher sensitivity, MetaPhlAn4 higher specificity, and Kraken2 lower overall performance.

These results demonstrate clear precision-recall trade-offs in metagenomic profiling. This benchmarking framework provides a reproducible approach for evaluating pipeline performance across diverse microbiome contexts and life stages.

## Introduction

The human microbiome is increasingly considered a key correlate of health. Microbiome profiles are associated with immune development, metabolic function, and disease risk (Samarra et al., 2025; Van Rossum et al., 2020). As a result, microbiome sequencing data have become a foundational input for basic research, translational studies, and emerging clinical applications. However, the accuracy of these insights depends fundamentally on the performance of the computational pipelines used to analyze metagenomic sequencing data. Methodological choices made during sequencing and analysis shape reported microbiome profiles, even when the underlying biological material is identical. Comparative studies have shown that differences in sequencing platform, library preparation, and bioinformatic pipeline can shift taxonomic composition and diversity estimates, which in turn can result in differing clinical interpretations (Allali et al., 2017; Servetas et al., 2026). These challenges can be especially consequential in early life, when the microbiome is highly dynamic, so even minor analytical differences may influence how developmental changes are interpreted (Nunez et al., 2025).

Taxonomic profiling faces technical challenges due to the complexity and diversity of microbial communities. Real-world microbiomes are highly complex, containing a few abundant taxa alongside many rare species that may still be biologically significant (Jousset et al., 2017). Additionally, microbial species often include diverse strains with varying genomes, which can affect how metagenomic pipelines assign reads and estimate abundances (Truong et al., 2017).

Different computational approaches employ distinct strategies for taxonomic assignment, each with specific strengths and limitations that are influenced in part by the reference genome database employed. Marker gene-based methods like MetaPhlAn4 identify organisms by detecting clade-specific marker genes within sequencing reads (Blanco-Míguez et al., 2023). In contrast, k-mer-based classifiers like Kraken2 assign taxonomy by matching short DNA sequences (k-mers) from reads directly to reference genomes (Wood et al., 2019). These methods must balance precision (correctly assigning sequences to taxa) with recall (detecting all taxa truly present), a trade-off that becomes particularly pronounced when identifying low-abundance species in complex communities. Despite the proliferation of metagenomic analysis pipelines, robust validation of their performance remains challenging. Because popular pipelines including MetaPhlAn4, Kraken2, mOTU, and Kaĳu, among others, each employ different computational strategies and reference databases, their outputs and performance can differ meaningfully, underscoring the need for careful benchmarking (Edwin et al., 2024; Odom et al., 2023; Parks et al., 2021; Poussin et al., 2022).

Evaluating metagenomic pipelines requires consideration of both community-level and taxon-specific performance. Global metrics such as precision, recall, F1 score, and AUPR are commonly used to quantify how accurately pipelines reconstruct the overall taxonomic composition of a microbial community. For example, AUPR evaluates performance across all detection thresholds and reflects how well a pipeline balances identification of both abundant and rare taxa while controlling false positives, an essential feature for capturing overall performance in complex microbial communities. However, global metrics can mask errors affecting individual taxa that are critical for biological interpretation. Pipelines differ in how they assign reads and balance precision and recall. As a result, two methods with similar overall metrics may perform very differently for specific species, potentially leading to missed or incorrectly identified key taxa. High performance at both the community and species level is essential to capture the full complexity of the microbiome and to accurately interpret the functional roles of individual taxa and their contributions to host biology.

Benchmarking is typically done with ground-truth datasets, which consist of samples with a precisely known composition of microbial species. While existing benchmarking approaches provide valuable insights, they have several important limitations when applied to real-world human microbiome samples.

1) Commercial reference communities provide a controlled and reproducible way to benchmark analytical accuracy, but their simplified microbial composition makes them less representative of real human samples. 2) Synthetic mock communities can overrepresent common microbiome configurations, limiting assessment of pipeline performance across the full diversity of microbial community types observed in human populations. 3) Methods optimized for adult samples often generalize poorly across life stages, particularly for infant microbiomes, which differ substantially in composition and developmental dynamics (Beller et al., 2021; Bokulich et al., 2020; Milani et al., 2017). 4) Different body sites present unique analytical challenges, such as the low-diversity, *Lactobacillus*-dominated communities commonly found in the vaginal microbiome.

To address these gaps, we developed a comprehensive framework for generating synthetic metagenomes that capture the natural diversity of human microbiomes across sample types, including age stratification for gut samples to reflect key developmental life stages. To reduce redundancy and better capture microbiome diversity, we applied novelty-based sampling to select representative samples from adult gut, adult vaginal, child gut, toddler gut, and infant gut groups, prioritizing taxonomic dissimilarity to include both common and rare community types. We then constructed synthetic metagenomes by integrating multiple strains per species from both internal and external reference databases, simulating realistic sequencing conditions while maintaining known ground truth. Previous benchmarking efforts suggest that k-mer based pipelines are generally more comprehensive and faster, while marker-gene methods provide more precise measurements for certain taxa (Edwin et al., 2024; Miossec et al., 2020; Poussin et al., 2022). Our approach enabled systematic validation of three metagenomic pipelines: Tiny Health’s proprietary Metagenomic Classifier v2 (THMCv2), MetaPhlAn4, and Kraken2+Bracken, evaluating their ability to accurately identify taxa across multiple age groups and sample types.

This study addresses a critical need in the field by providing a framework for rigorous taxonomic validation across microbiome contexts. By benchmarking pipeline performance on age-specific synthetic communities that reflect real-world diversity, we quantify the precision-recall trade-offs inherent in metagenomic profiling and assess each method’s ability to detect both abundant and rare taxa. We further evaluate species-level sensitivity and specificity for clinically relevant taxa, including infant gut members, opportunistic pathogens, commensal adult microbes, and vaginal community indicators, to assess performance in contexts where misclassification carries direct interpretation consequences.

## Methods

### Initial sample processing

To create synthetic metagenome mock communities with a known ground truth that ensure broad representation of microbiomes across different age groups and sampling types, 5,000 samples were randomly chosen from Tiny Health reference data for each category: adult gut samples (>18 years of age), child gut samples (3-18 years of age), toddler gut samples (1-3 years of age), infant gut samples (<1 year of age, further divided into >6 months and <6 months), and adult vaginal samples.

Samples were collected, processed, sequenced, and taxonomically profiled as previously described in (Nieto et al., 2025). Briefly, samples were collected at home using FLOQSwab-ADT collection kits (Copan, USA) and processed in a CLIA/CAP-certified laboratory. Total DNA was extracted using the DNeasy PowerSoil Pro kit (Qiagen, Germany), and sequencing libraries were prepared using the Nextera kit (Illumina, USA). Samples were then sequenced on an Illumina NovaSeq platform (Illumina, USA) to a minimum depth of 10 million reads.

Raw reads underwent quality control using Fastp (Chen et al., 2018), including adapter and low-quality sequence trimming, duplicate read removal, and filtering of human-derived reads. The remaining high-quality non-host reads were used for species-level taxonomic classification against a comprehensive microbial reference database spanning bacterial and archaeal genomes from GTDBv202 (Parks et al., 2022), and viral, fungal, and protistan genomes available through NCBI GenBank (Sayers et al., 2025).

### Ground truth community selection

From the initial pool of 5,000 samples per category, a systematic selection process was used to identify 50 samples per category encompassing a wide range of microbiome compositions. This process aimed to select a set of samples that covered as much taxonomic diversity as possible and proceeded as follows: 1) Pairwise Bray-Curtis distances were calculated between all samples; 2) A random starting sample was chosen to serve as the seed for the pool of selected samples; 3) For each sample in the unselected pool, a novelty score was calculated as the mean distance to already selected samples multiplied by the minimum distance to already selected samples; 4) The sample with the highest novelty score was selected, added to the selected sample pool, and removed from the unselected sample pool. Steps 3 and 4 were repeated until each category was represented by 50 samples.

### Synthetic metagenome generation

To construct representative synthetic metagenomes, species-level taxonomic profiles were retrieved for each chosen sample from the Tiny Health database. The relative abundance of each species in the ground truth samples was derived from these taxonomic profiles. Only species with a relative abundance above 0.005% were included, which aligns with expectations for reliable detection of low abundance features in this field (Poussin et al., 2022). To create a synthetic sample, genomes that were publicly available were selected to match the taxonomic profiles of the original samples. All genomes present were downloaded using NCBI datasets and data format (O’Leary et al., 2024). Because real world samples could be derived from multiple strains of the same species (Van Rossum et al., 2020), strain-level diversity was simulated by choosing 1-4 genomes for every species present, each corresponding to a distinct strain. When multiple strains per species were present, the relative abundance of that species was equally divided among the strains.

Because real-world samples will not be made of reference genomes, genomes outside of the Tiny Health taxonomic reference database were prioritized, although some genomes present in the Tiny Health taxonomic reference database were included when the number of available representative genomes for that species was limited. Using this genome-level ground truth abundance data, synthetic metagenomes were generated. BBtools randomreads.sh was used to generate 10,000,000 random synthetic Illumina-like 2x150 PE reads (5,000,000 read-pairs) using the downloaded genomes and their corresponding relative abundances in the ground truth data.

### Taxonomic Classification

Three different taxonomic classifiers were compared: the Tiny Health proprietary pipeline (THMCv2), MetaPhlAn4, and Kraken2+Bracken with a standard NCBI taxonomy-based reference database (herein referred to as Kraken2). THMCv2 uses the GTDB taxonomy r202 release. MetaPhlAn4 version 4.1.0 was used with the mpa_vJun23_CHOCOPhlAnSGB_202307 SGB database. Conversions between the SGB taxonomy and the GTDB taxonomy were performed using the file mpa_vJun23_CHOCOPhlAnSGB_202307_SGB2GTDB.tsv, which is provided with MetaPhlAn4. Kraken2+Bracken was run using a “standard” NCBI-based database built on January 12th, 2024 and provided via https://benlangmead.github.io/aws-indexes/k2. All tools were run with default settings.

### Calculating validation metrics

To evaluate taxonomic classification performance, four standard metrics were used: precision, recall, F1 score, and area under the precision-recall curve (AUPR). For all metrics, true positives (TP), false positives (FP), false negatives (FN), and true negatives (TN) were calculated. Precision, calculated as TP / (TP + FP), is defined as the proportion of correctly assigned sequences among all assignments. Recall, calculated as TP / (TP + FN), is defined as the proportion of true taxa that are correctly detected. The F1 score, calculated as 2 × (Precision × Recall) / (Precision + Recall), is defined as a balance between precision and recall into a single score. The AUPR summarizes classifier performance across all thresholds, capturing its ability to detect both abundant and rare taxa while minimizing incorrect assignments. In addition to these standard metrics, which generally do not take the abundance of taxonomic classifications into consideration, “weighted” metrics were also calculated, where the standard metrics were weighted by the relative abundance of each feature. Abundance-weighted precision was calculated via sum(TP) / (sum(TP) + sum(FP)). Abundance-weighted recall was calculated via sum(TP) / (sum(TP) + sum(FN)). The average abundance of FP and FN classifications was also calculated for each pipeline.

In addition to these community-wide validation metrics, sensitivity and specificity were also reported for select taxa. Sensitivity is equivalent to recall, while specificity is calculated via TN / (TN + FP). The relative abundance and prevalence of each selected taxon were also calculated.

### Statistics

To compare the validation metrics between taxonomic classifiers, linear models were used and contrasts of interest extracted with the R package emmeans (Lenth & Piaskowski, 2026), with p-values adjusted for multiple comparisons using Tukey’s HSD.

## Results

### Mock communities represent diverse sample types

Novelty-based sampling was used to construct the ground truth communities, prioritizing representative coverage of diverse microbiomes (**Figure 1**). Relative abundance profiles of the top 16 most abundant bacterial families were then examined across age groups and sampling types using stacked bar plots (**Figure 2**). Vaginal microbiomes were characterized by prominent *Lactobacillaceae*, *Bifidobacteriaceae*, and *Bacteroidaceae* (primarily *Prevotella spp*.), distinguishing them from gut communities (**Figure 2**). Adult, child, toddler, and infant samples older than 6 months were dominated primarily by *Lachnospiraceae* and *Bacteroidaceae*, with *Lachnospiraceae* lowest in infants and generally increasing with age (**Figure 2**). Additionally, infants under 6 months showed more variable family-level profiles, with some samples enriched for *Enterobacteriaceae* or *Bifidobacteriaceae* (**Figure 2**). Overall, these patterns highlight intrinsic variability across microbiomes and reveal distinct shifts in composition across sample types and developmental stages.

**Figure 1:**
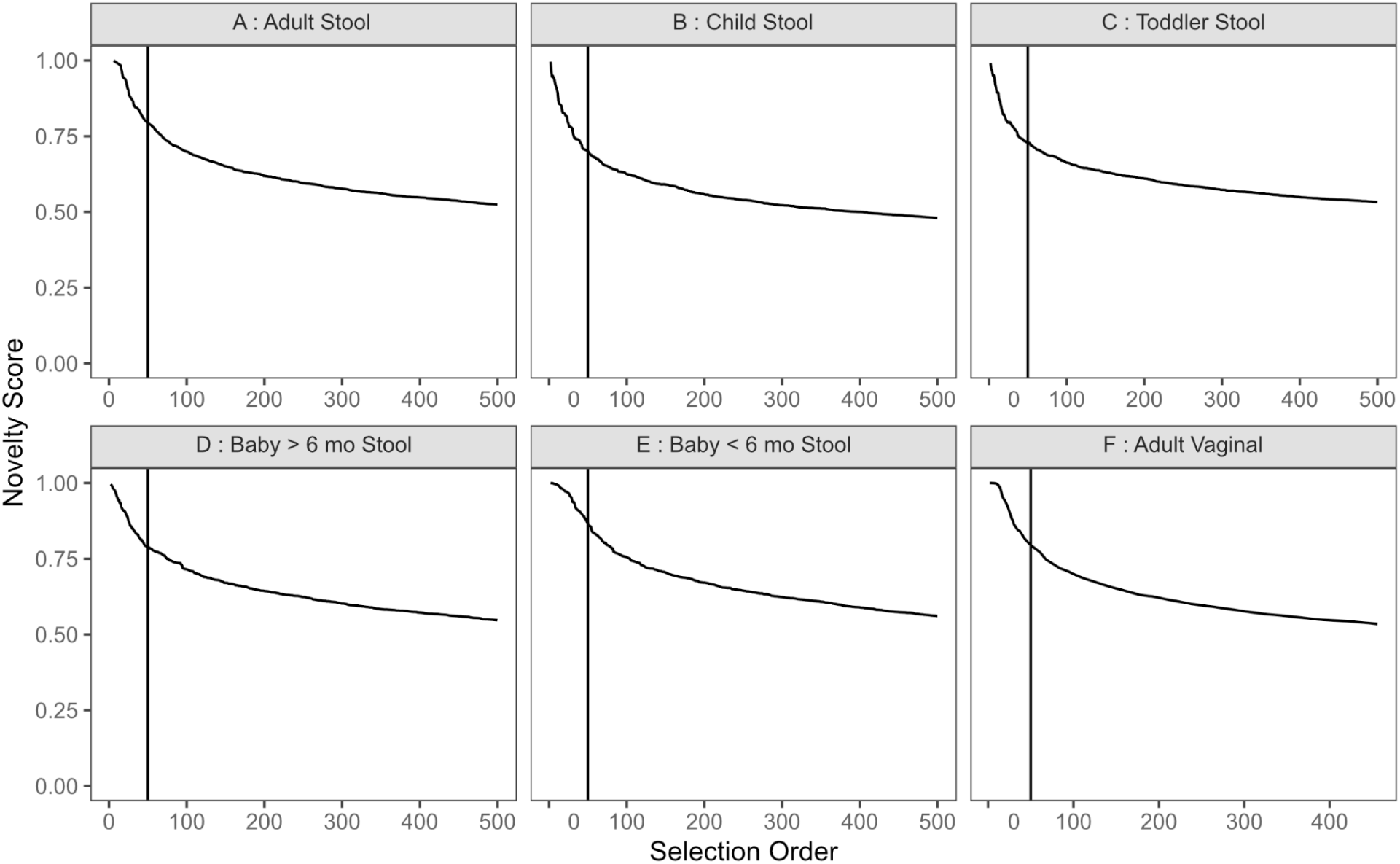
Novelty of selected samples decreased rapidly after selection of the first 50 samples. Novelty score curves for samples from each of the age and body-site categories. The x-axis indicates selection order and the y-axis indicates novelty score. The vertical line marks the 50th sample included in the final selected sample pool.

**Figure 2:**
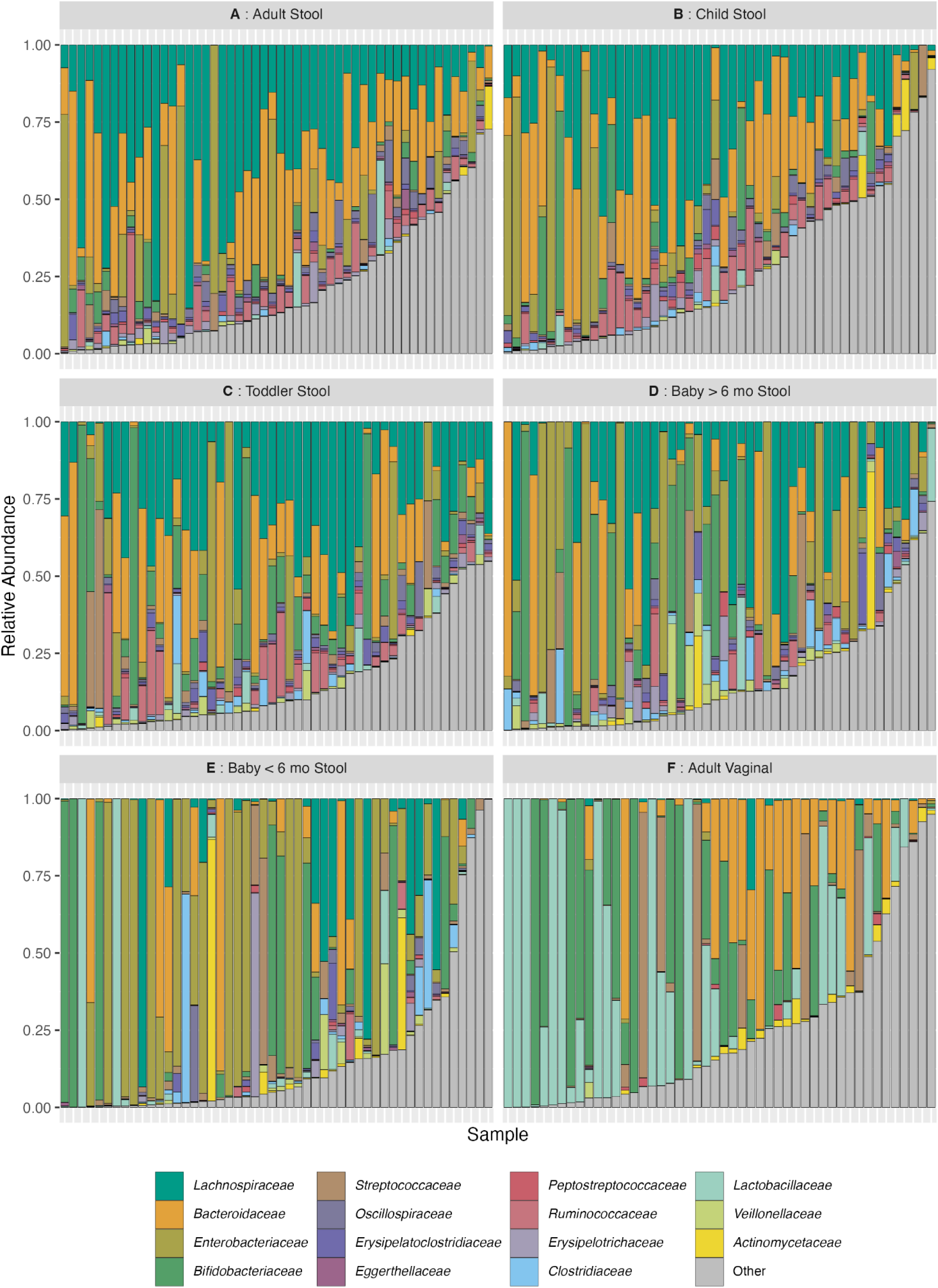
Microbiome composition differs across age and body-site. Stacked bar plots depicting the abundance of the top 16 families across samples used as templates for the synthetic metagenomes.

### Synthetic metagenome communities are primarily generated from outside reference databases

To construct representative synthetic metagenomes, taxonomic profiles were retrieved for each chosen sample. To realistically simulate real-world conditions, we preferentially selected genomes from outside of the Tiny Health reference database. While numerically most strains were sourced from within our reference database, the majority of the microbiome community, as measured by the relative abundance, was sourced from outside the reference database in most age categories (**Figure 3**).

**Figure 3:**
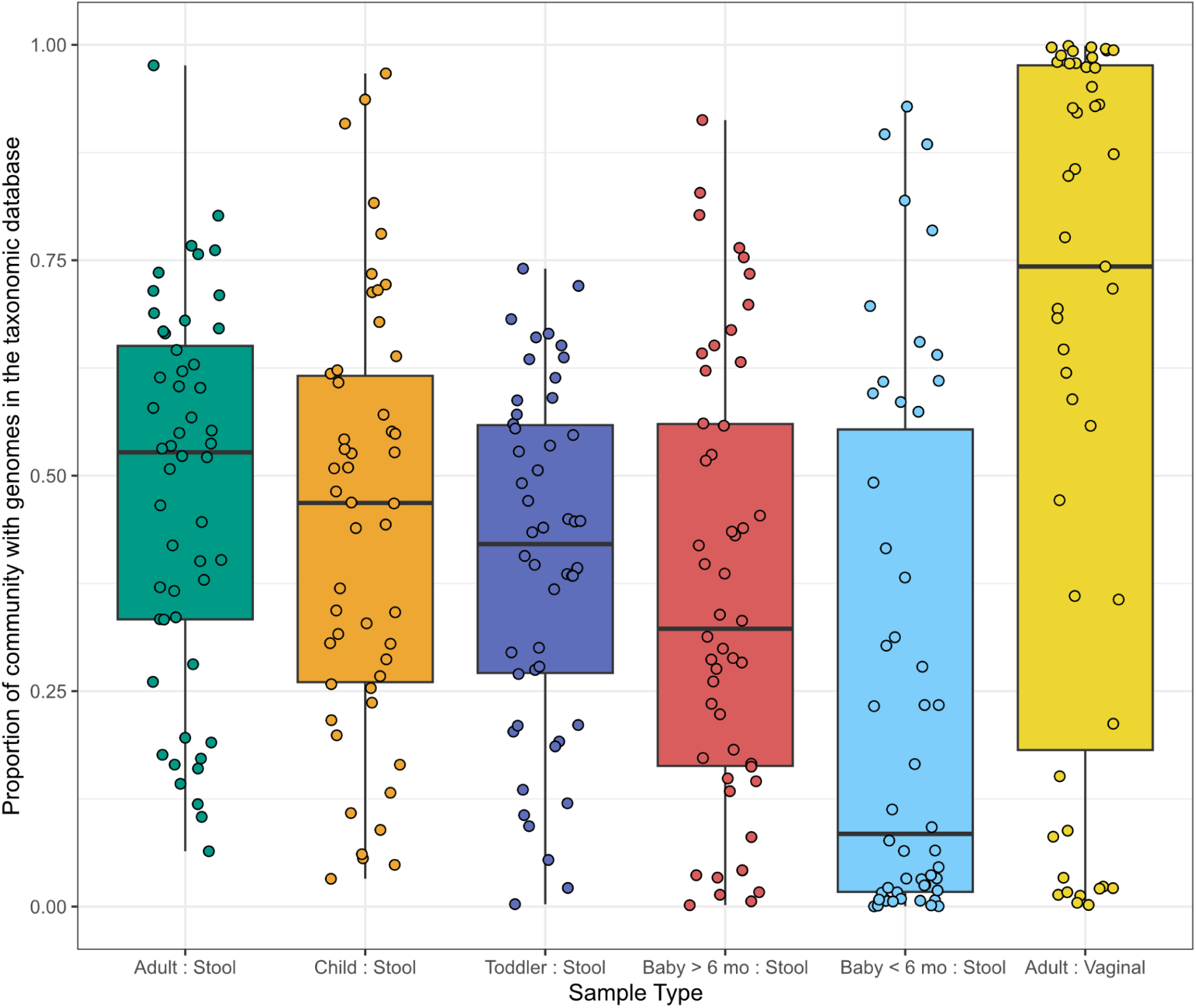
Most of the community abundance was derived from strains outside the Tiny Health reference database. The plotted values represent the relative abundance from strains with genomes in the database. Boxplots show the median, interquartile range, and individual data points for each age- and body-site category.

### Classification accuracy is highest for THMCv2, followed by MetaPhlAn4, then Kraken2

To measure the performance of taxonomic classifications, four standard evaluation metrics were utilized: precision, recall, F1 score, and area under the precision-recall curve (AUPR). The results of this validation framework showed that both THMCv2 and MetaPhlAn4 performed well with each exhibiting strengths in different performance metrics while Kraken2 using the standard DB exhibited lower performance. All comparisons were statistically significant with an adjusted p-value < 0.05. Full pairwise comparison statistics are provided in **Supplementary Table S1**.

Across all ages and sample types, THMCv2 showed substantially higher recall than both MetaPhlAn4 and Kraken2, indicating a stronger ability to recover taxa truly present in the samples (Tukey’s HSD, p < 0.01**) (Figure 4)**. In contrast, MetaPhlAn4 achieved the highest precision across all ages and sample types (Tukey’s HSD, p < 0.05) (**Figure 4**), while Kraken2 showed the lowest precision, reflecting a higher rate of false-positive taxonomic assignments (Tukey’s HSD, p < 0.0001). The difference between MetaPhlAn4 and THMCv2 was most pronounced in infant samples (Tukey’s HSD, p < 0.0001) (**Figure 4**).

**Figure 4:**
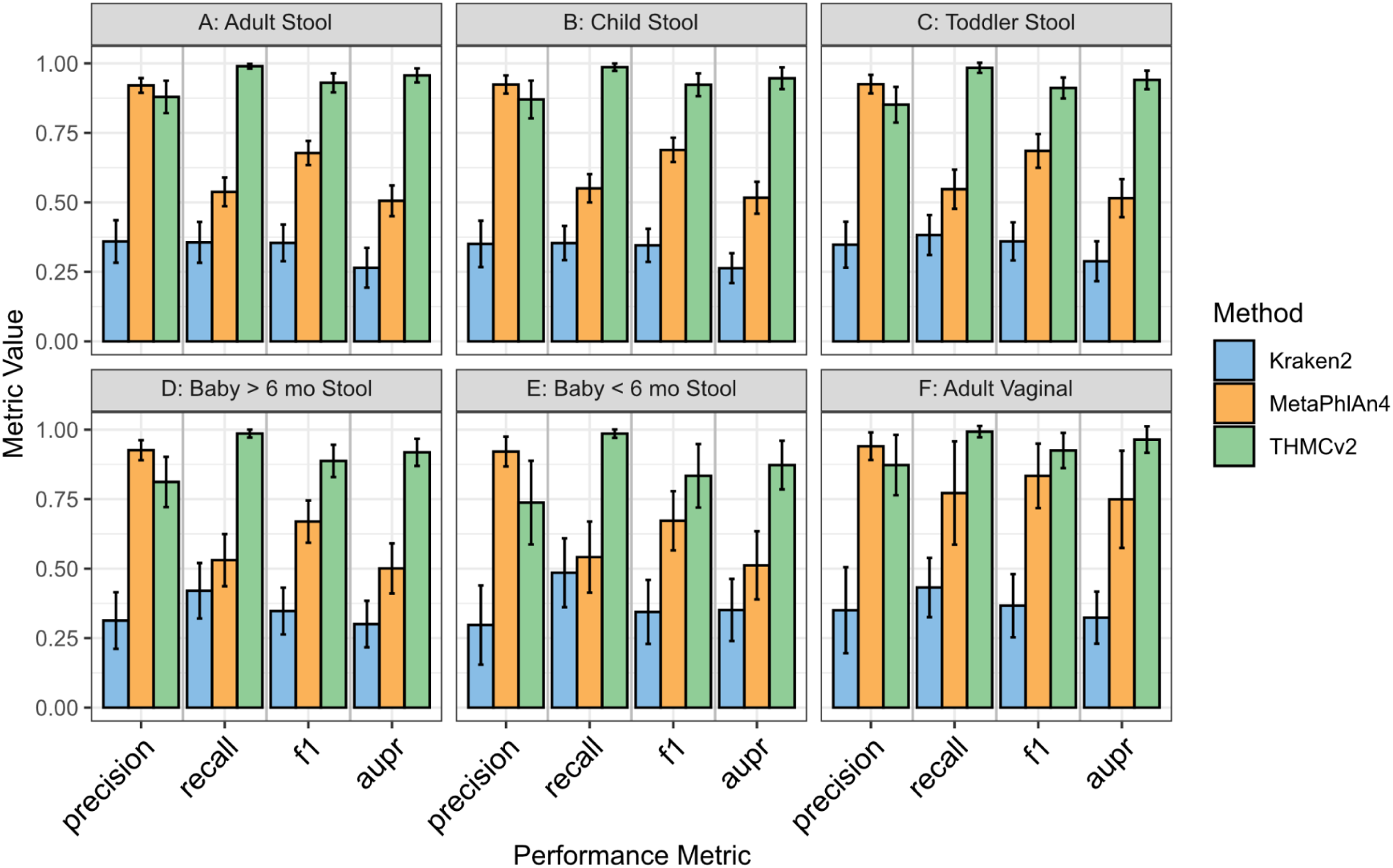
Classification accuracy is highest for THMCv2, followed by MetaPhlAn4, then Kraken2. Barplots represent precision, recall, F1, and AUPR values for the three analysis pipelines across age and body-site categories: (A) adult stool, (B) child stool, (C) toddler stool, (D) baby >6 mo stool, (E) baby <6 mo stool, and (F) adult vaginal. Error bars represent the mean +- 1 standard deviation. Statistical differences between pipelines were assessed using a linear model with Tukey’s HSD post hoc tests (p < 0.05), and all comparisons were statistically significant.

When precision and recall were considered together, THMCv2 achieved the highest F1 scores across all sample types and age groups, reflecting the best overall balance between these metrics among the methods tested (Tukey’s HSD, p < 0.0001) **(Figure 4)**. Similarly, THMCv2 achieved the highest AUPR across all sample types and ages (Tukey’s HSD, p < 0.0001) **(Figure 4)**. AUPR is valuable for metagenomics because it measures how well a classifier balances true positive and false positives in datasets dominated by a few abundant and many rare species. Unlike metrics that emphasize true negatives, AUPR focuses on this precision-recall trade-off, making it especially relevant for accurately identifying taxa in highly imbalanced communities.

### Abundance-weighted performance is comparable between THMCv2 and MetaPhlAn4

A typical microbiome comprises both highly abundant and extremely rare taxa. Although species with less than a 0.005% relative abundance were excluded during initial data processing, in this analysis results were adjusted to account for each species’ relative abundance. This approach highlights the ability of pipelines to identify dominant taxa and reduces the influence of low-abundance taxa on performance metrics. Full pairwise comparison statistics are provided in **Supplementary Table S2**.

Across each measure, THMCv2 and MetaPhlAn4 achieved nearly perfect performance when weighted by abundance, while Kraken2 showed lower scores in each case (Tukey’s HSD, p < 0.01) (**Figure 5**). However, the focus on highly abundant species in this analysis inflates the performance of each pipeline, as it softens the impact of missing or incorrectly identifying rare species.

**Figure 5:**
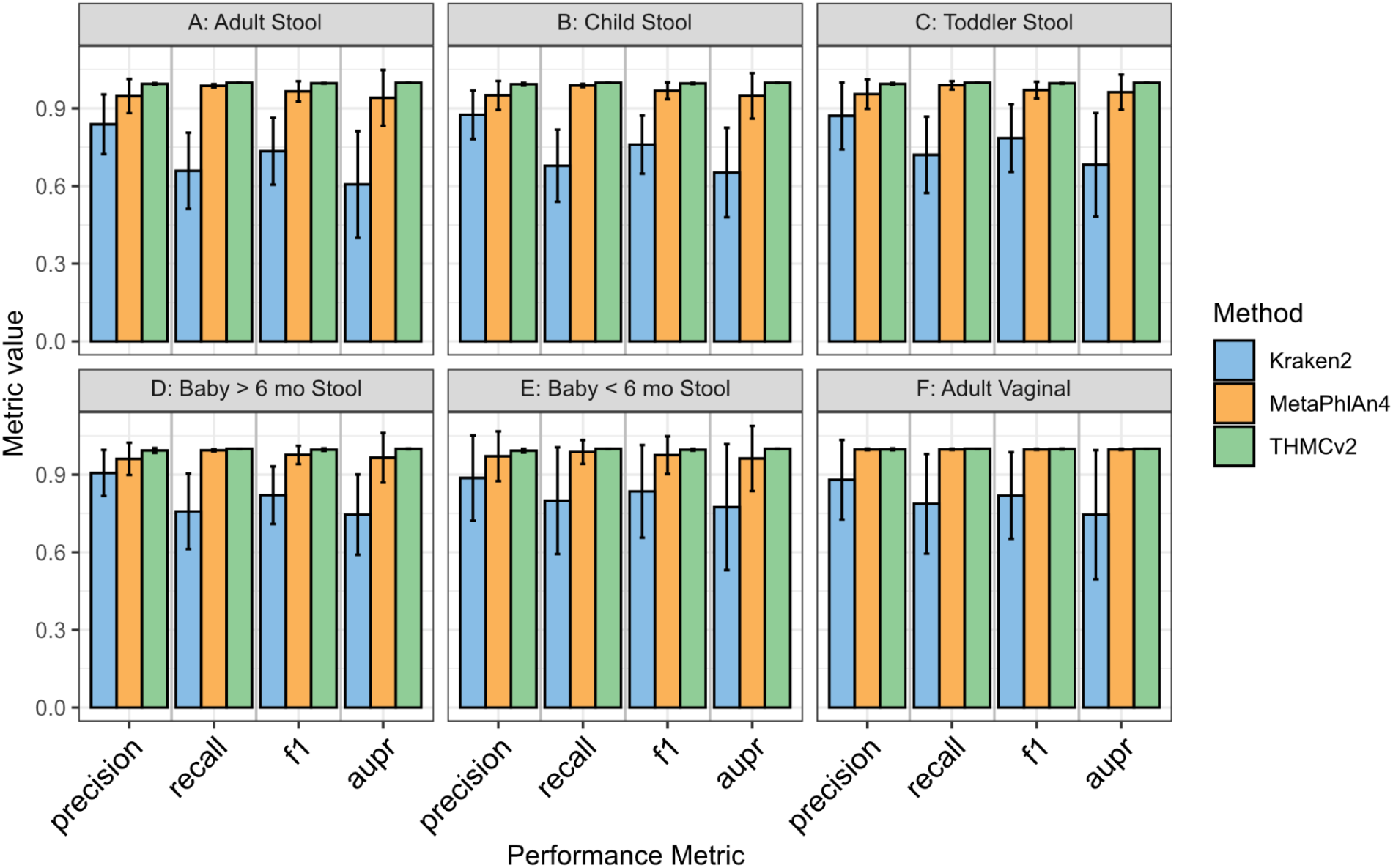
THMCv2 and MetaPhlAn4 have comparable abundance-weighted classification accuracy, while Kraken2 performs worse. Barplots represent abundance-weighted precision, recall, F1, and AUPR values for the three analysis pipelines across age and body-site categories: (A) adult stool, (B) child stool, (C) toddler stool, (D) baby >6 mo stool, (E) baby <6 mo stool, and (F) adult vaginal. Error bars represent the mean +- 1 standard deviation. Statistical differences between pipelines were assessed using a linear model with Tukey’s HSD post hoc tests (p < 0.05).

We next investigated the abundance of features subject to classification errors. Note that this is not a measurement of the number of false positives or negatives, but rather the abundance of features that belong to these error categories. Full pairwise comparison statistics are provided in **Supplementary Table S3**. Across all three tools, false negatives were generally low abundance features (low abundance in the ground truth communities). Across all ages, false negatives from THMCv2 and MetaPhlAn4 exhibited lower abundance relative to those by the Kraken2 classifier (p < 0.0001) (**Figure 6**). In vaginal samples, false negatives by the MetaPhlAn4 classifier were of lower abundance than Kraken2 only (p < 0.05) (**Figure 6**). This indicates that when these pipelines make a false negative type error (i.e. misses a taxa), this taxa is likely a very low abundance member of the community.

**Figure 6:**
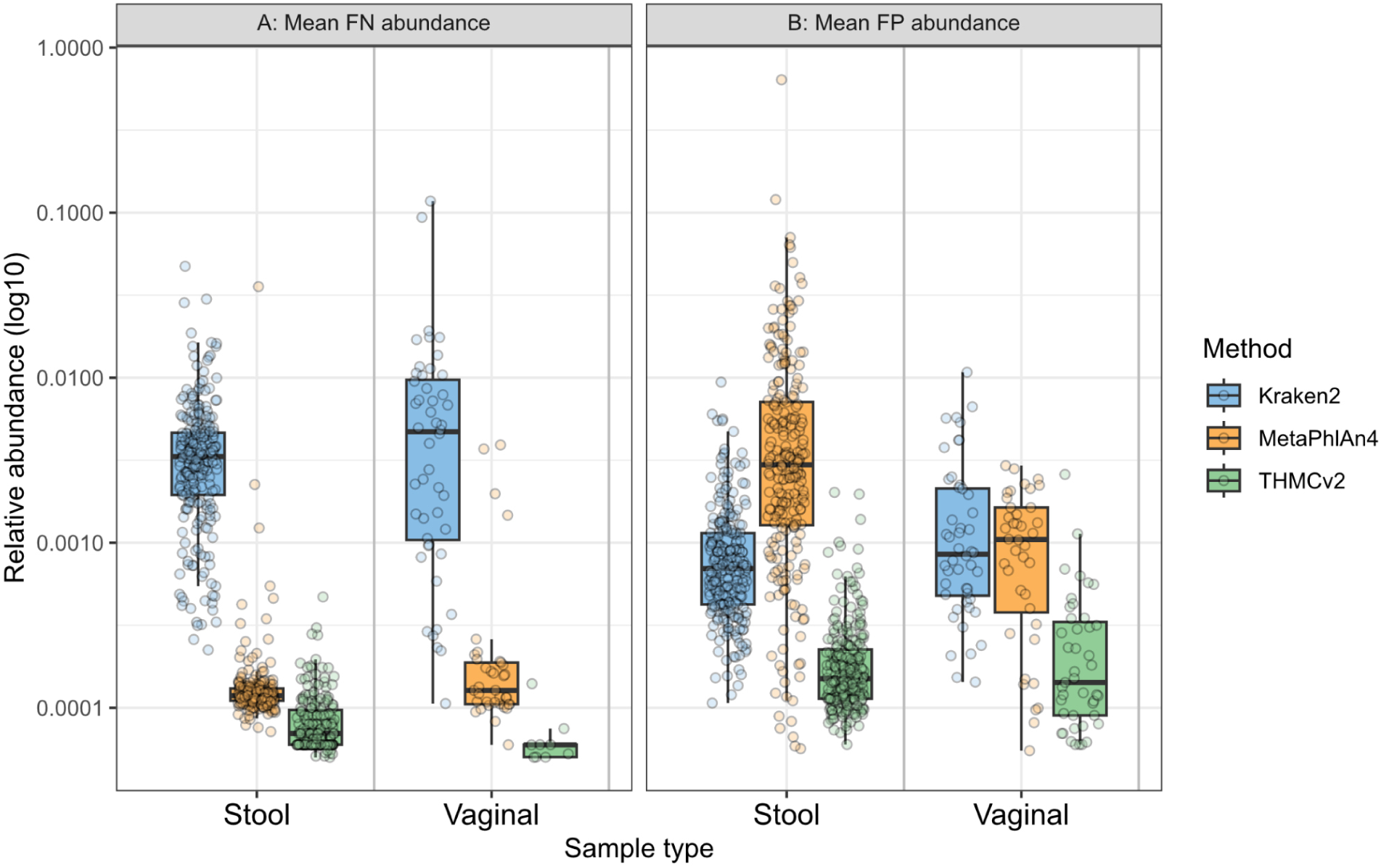
For THMCv2, classification errors were generally only found among low abundance taxa. Boxplots represent the average abundance of (A) False Negative (FN) classifications and (B) False Positive (FP) classifications in stool and vaginal samples, for each pipeline. It is important to note that these boxplots do not depict the number of such events, but rather, when these events occurred, what the abundances of these features were. For the FN events, the ground truth abundance is plotted, for the FP events, the abundance reported by the classifier is plotted.

We similarly investigated the abundance of features identified in false positive type errors (i.e. incorrectly identifying an absent species). Across all ages, with the exception of infants under 6 months of age, the THMCv2 and Kraken2 pipelines produced false positives of lower abundance than false positives from MetaPhlAn4 (p < 0.0001) (**Figure 6; Supplementary Table S3**). In infants under 6 months of age the abundance of features identified in false positive errors was equal across pipelines (p > 0.05). Notably, despite having the highest unweighted precision, MetaPhlAn4 false positives tended to be of higher abundance than those from the other pipelines. Overall, errors from THMCv2, both false positives and false negatives, were generally of extremely low abundance.

### Species-specific performance varies across pipelines

While aggregate metrics provide an overall assessment of pipeline performance, interpretable microbiome analysis requires accurate identification of specific microbial taxa. We therefore evaluated pipeline performance at the species level using sensitivity and specificity. Sensitivity reflects a pipeline’s ability to detect a species when present, directly minimizing false negatives, a critical consideration for identifying potential pathogens or beneficial commensals that inform actionable decisions. Specificity measures a pipeline’s ability to correctly identify when a species is absent, minimizing false positives that could lead to erroneous biological interpretations.

We stratified our species-level analysis into four categories that represent distinct validation challenges: key infant gut members, clinically relevant opportunistic pathogens, common adult microbiome members, and vaginal community state type (CST) indicator species. This approach allowed us to assess not only overall pipeline accuracy but also performance in contexts where misclassification carries interpretation consequences.

Analysis of infant gut members revealed the importance of accurate early-life microbiome profiling. We assessed the ability to identify several highly developmentally-relevant taxa such as the *Bifidobacterium* species, *B. longum*, *B. breve, B. bifidum*, and *B. animalis*, as well as *Lacticaseibacillus rhamnosus*. High specificity was achieved for *B. bifidum*, *B. animalis,* and *L. rhamnosus* by all three pipelines. However, specificity for *B. longum* and *B. breve* varied widely across pipelines, with exceptional performance by MetaPhlAn4, but lower specificity for *B. longum* by THMCv2 and for both *B. longum* and *B. breve* by the Kraken2 pipeline (**Figure 7A**). In contrast, THMCv2 showed consistently high sensitivity across all *Bifidobacterium* species and *L. rhamnosus*, with variable performance by both Kraken2 and MetaPhlAn4 (**Figure 7B**).

**Figure 7:**
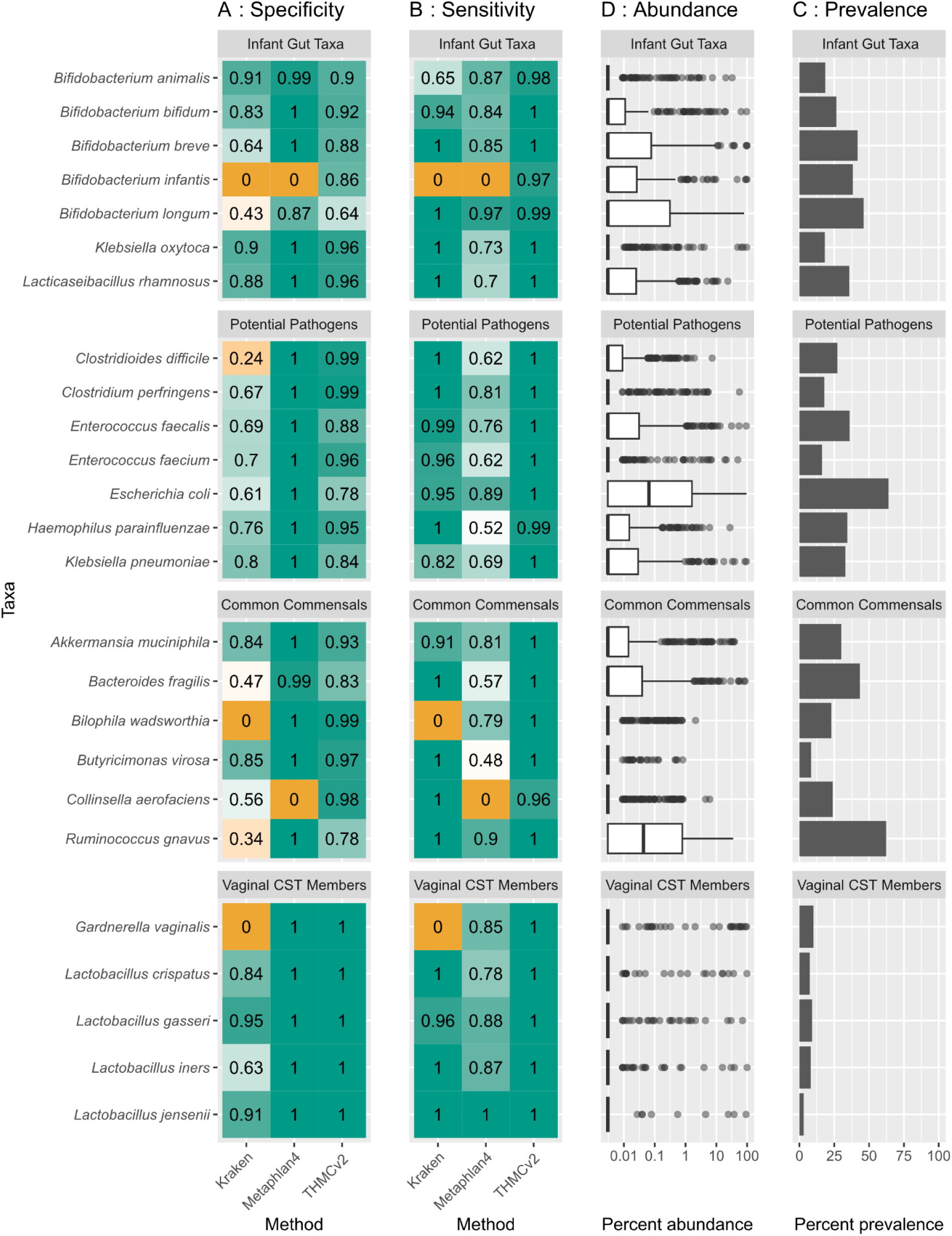
THMCv2 achieves the best balance of species-level specificity and sensitivity compared with MetaPhlAn4 and Kraken2. (A) Specificity and (B) sensitivity values for infant gut taxa, opportunistic pathogens, common gut commensals, and vaginal CST indicator species. Color shading ranges from orange to green (0 to 1), with green indicating higher specificity and sensitivity scores. (C) Relative abundance and (D) prevalence of each species within these groups in all sample types, with CST indicator species in vaginal samples only.

We also assessed the identification of a potentially disruptive species present in infant gut microbiomes, *Klebsiella oxytoca*. Specificity was high for this species in all three pipelines (**Figure 7A**). Kraken2 and THMCv2 identified this species with perfect sensitivity, while MetaPhlAn4’s sensitivity was lower (**Figure 7B**).

We next assessed the correct identification of potential pathogens of interest, including *Clostridioides difficile, Clostridium perfringens, Enterococcus faecalis, Enterococcus faecium*, *Escherichia coli, Haemophilus parainfluenzae,* and *Klebsiella pneumoniae.* The pipelines achieved similar specificity and sensitivity across these potential pathogens. Specificity was highest for MetaPhlAn4, intermediate for THMCv2, and lowest for Kraken2 (**Figure 7A**). In contrast, sensitivity was highest for THMCv2, with intermediate performance by Kraken2, and the lowest sensitivity by MetaPhlAn4 (**Figure 7B**).

Next, we assessed the accurate detection of common commensal adult species. For *Akkermansia muciniphila, Bacteroides fragilis, Bilophila wadsworthia, Butyricimonas virosa, Collinsella aerofaciens,* and *Ruminococcus gnavus,* specificity was highest for MetaPhlAn4, with slightly lower specificity by THMCv2. Specificity was much lower for Kraken2, suggesting a failure to minimize false positives (**Figure 7A**). Sensitivity was nearly perfect across this group of species by both Kraken2 and THMCv2, but much lower for MetaPhlAn4 (**Figure 7B**).

In vaginal samples, accurate identification of CST indicator species is essential for understanding microbiome health. We assessed the identification of *Lactobacillus crispatus*, *Lactobacillus gasseri*, *Lactobacillus iners*, and *Lactobacillus jensenii*, hallmarks of CSTs 1, 2, 3, and 5, respectively, and *Gardnerella vaginalis*, a common species in CST 4. Specificity was perfect for both MetaPhlAn4 and THMCv2, and moderate performance was also seen for Kraken2 for all species other than *G. vaginalis* (**Figure 7A**). Sensitivity was perfect for THMCv2 and near-perfect for Kraken2, with MetaPhlAn4 also performing well for all species other than *G. vaginalis* (**Figure 7B**).

Relative abundance and taxa prevalence do not explain the observed differences in pipeline performance. Several taxa with low abundance, including *Bifidobacterium bifidum, Akkermansia muciniphila*, and vaginal CST species such as *Lactobacillus gasseri* and *L. jensenii*, were recovered with high sensitivity and specificity across pipelines (**Figure 7C, D**). Notably, near-perfect classification was achieved for *L. jensenii* across pipelines, despite relatively low prevalence (**Figure 7D**). In contrast, taxa with broader abundance distributions, including *Bifidobacterium longum* and *Escherichia coli*, showed pipeline-dependent variation in specificity (**Figure 7C**), and this was not resolved by prevalence: *E. coli* is among the most prevalent taxa examined, yet still exhibited inconsistent specificity across pipelines (**Figure 7D**). Likewise, some low-abundance, low-prevalence taxa like *Collinsella aerofaciens* were well resolved by at least one pipeline (**Figure 7C, D**). Overall, these results indicate that intrinsic characteristics of taxa, along with pipeline-specific strategies, play major roles in shaping sensitivity and specificity across groups, more so than abundance or prevalence.

## Discussion

By applying an age- and body-site stratified synthetic metagenome framework, we explored how taxonomic classification accuracy varies between pipelines as well as across sample types and life stages. While MetaPhlAn4 often showed the highest precision, THMCv2 demonstrated superior recall and achieved the highest F1 scores and AUPR across all sample types, indicating the best overall balance between accuracy and comprehensiveness. When weighted by abundance, both THMCv2 and MetaPhlAn4 achieved near-perfect performance. When THMCv2 produced false positives and false negatives, they were of lower abundance (<0.001%), indicating that when misidentifications occur, they are predominantly limited to very low-abundance features.

Species-specific analyses allowed us to assess performance across a variety of biological contexts. Performance patterns were consistent across these contexts, with strong specificity by the THMCv2 and MetaPhlAn4 pipelines but lower specificity by Kraken2. Sensitivity was high for both THMCv2 and Kraken2 and much lower for MetaPhlAn4. Overall, THMCv2 maintained a balance between these measures, whereas Kraken2 produced more false positives and MetaPhlAn4 more false negatives.

Our results demonstrate that pipeline performance varies across age groups and sample types, underscoring the critical need for age-stratified validation in microbiome research. The compositional differences between infant, child, and adult microbiomes, as well as between gut and vaginal communities, present distinct analytical challenges that cannot be adequately captured by generalized or reference communities. For instance, MetaPhlAn4’s stronger precision in infant samples compared to other age groups suggests that marker gene-based approaches may be particularly well-suited to the less diverse taxonomic profiles characteristic of early life, where communities are often dominated by a smaller number of species from genera such as *Bifidobacterium (Milani et al., 2017)*. Conversely, the more complex, diverse adult gut microbiomes, with their greater abundance of *Bacteroidetes* and *Firmicutes* (Human Microbiome Project Consortium, 2012), may benefit from different pipeline strategies.

Further, the inclusion of vaginal samples in our validation framework addresses a typically underrepresented sample type in pipeline benchmarking. MetaPhlAn4 trended towards an improved performance in vaginal samples relative to gut samples, suggesting that its marker gene-based approach is similarly well suited to the low-diversity, vaginal microbiome, whereas it misses more taxa in the complex gut environment. We also assessed each pipeline’s ability to correctly identify vaginal CST indicators, and found that all three had strong specificity and sensitivity for the majority of these species. The ability to evaluate pipeline performance in this context ensures more comprehensive validation for applications in women’s health and reproductive medicine.

A central finding of our study is the fundamental trade-off between precision and recall in metagenomic taxonomic profiling. MetaPhlAn4 achieved the highest precision, particularly in infant samples, reflecting its conservative approach to taxonomic assignment through marker gene detection. This strategy minimizes false positives by requiring strong evidence before assigning sequences to specific taxa. However, this comes at the cost of recall, as MetaPhlAn4 consistently detected fewer true taxa than THMCv2 across all sample types and age groups.

In contrast, THMCv2 demonstrated substantially higher recall while maintaining competitive precision, resulting in higher F1 and AUPR values. This indicates a more aggressive detection strategy that successfully identifies a broader range of taxa, including low-abundance taxa that other methods may miss.

Our abundance-weighted analysis provides additional insight into this trade-off. When performance metrics were weighted by taxon abundance, all pipelines showed improved performance, and the gap between THMCv2 and the other methods diminished. This suggests that THMCv2’s advantage is particularly pronounced for low-abundance taxa, which comprise the majority of species-level features but a small fraction of total abundance. Consistent with this pattern, analysis of the abundance of false positives and false negatives shows that THMCv2’s misidentifications are concentrated in lower-abundance features, indicating that errors occur only when abundance drops to extremely low levels.

The practical implications of this trade-off depend on the research or clinical question at hand, as revealed by our species-level sensitivity and specificity analysis. For example, in the detection of clinically relevant opportunistic pathogens, THMCv2 achieved high specificity while maintaining near-perfect sensitivity, effectively balancing the need to detect organisms without producing false positives. Kraken2 performed with lower specificity, which indicates the presence of more false positives. The potentially more concerning scenario is in the case of MetaPhlAn4, which performed with lower sensitivity. This pipeline therefore produces more false negatives, which could potentially lead to missed bacterial species.

We found that variation in pipeline performance was not driven by taxon abundance or prevalence, as several low-abundance and low-prevalence taxa were recovered with high sensitivity and specificity across pipelines, while some highly abundant taxa showed pipeline-dependent variation. Instead, performance differences are more likely attributable to taxon-specific genomic characteristics and reference database representation. Marker gene-based methods like MetaPhlAn4 depend on the availability of unique clade-specific markers, which may not exist for all species or closely related taxa, while k-mer-based methods like Kraken2 can suffer reduced specificity when k-mers are shared across phylogenetically similar species (Blanco-Míguez et al., 2023; Wood et al., 2019).

While our study provides a framework for age- and body-site stratified pipeline validation, several limitations should be noted. First, although we prioritized strain genomes from outside the Tiny Health reference database to simulate realistic sequencing conditions, we were constrained to species already represented in our database for synthetic metagenome assembly. In real-world samples, previously uncharacterized species would likely be present, and metagenomic reads from these organisms could attempt to map to the most closely related species or strains in the reference database (Sun et al., 2026). This scenario would likely increase false-positive rates across all pipelines, as sequences from unknown taxa are incorrectly assigned to known, closely related species. Our validation may therefore underestimate the false-positive burden that pipelines encounter in practice, particularly in understudied populations or body sites where microbial diversity isn’t completely understood. Future work incorporating synthetic reads from organisms entirely absent from reference databases would provide a more stringent test of pipeline specificity.

Second, our ground truth samples do not include low-abundance species below 0.005% relative abundance. While justified by accepted limits of detection for shotgun metagenomics, real human microbiome samples could contain biologically relevant species present at these very low abundances. In some contexts, even extremely rare taxa can play important roles in community function or serve as early indicators of dysbiosis. Including ultra-low-abundance taxa in our validation would have markedly increased methodological complexity and introduced the risk of false signals from sequencing errors or contaminant reads. While our cutoff of 0.005% relative abundance may exclude relevant taxa, it is well below the 1% threshold below which a prior benchmarking study showed taxonomic pipelines often struggle to reliably detect taxa (Poussin et al., 2022).

Finally, it is important to note that the ground truth samples were created using taxonomy from GTDB, and while THMCv2 also uses GTDB, both Kraken2 and MetaPhlAn4 use alternative taxonomies. This necessitated a translation step between taxonomies for these two pipelines, which may have introduced an additional source of error into their results.

## Conclusion

We have developed an age-stratified framework for validating metagenomic profiling pipelines that captures the natural diversity of human microbiomes across critical life stages and body sites. Applying this framework, we identified clear differences in pipeline performance, with each method showing unique strengths and weaknesses in precision and recall, especially for detecting low-abundance and clinically relevant taxa. Our findings underscore the necessity of rigorous, context-aware benchmarking to accurately assess pipeline capabilities, guiding the informed selection and improvement of metagenomic tools. This approach enhances reproducibility in microbiome research and supports the development of reliable pipelines essential for translating metagenomic data into meaningful applications.

