## Supplemental Tables 1-3 for "Robust taxonomic classification in gut and vaginal microbiomes demonstrated through benchmarking with age-specific synthetic communities"

**Table S1.** Pairwise comparisons of classification performance metrics across methods, grouped by body site and age group. Estimates reflect mean differences between methods. Adjusted p-values were corrected for multiple comparisons. SE, standard error; CI, confidence interval; df, degrees of freedom.

| Metric | Group | Contrast | Estimate | SE | df | 95% CI low | 95% CI high | t | adj. p |
|---|---|---|---|---|---|---|---|---|---|
| precision | Stool Adult | MetaPhlAn4 - Kraken2 | 0.5615 | 0.0113 | 153 | 0.5347 | 0.5882 | 49.658 | < 0.0001 |
| precision | | THMCv2 - Kraken2 | 0.5201 | 0.0113 | 153 | 0.4934 | 0.5469 | 46.001 | < 0.0001 |
| precision | | THMCv2 - MetaPhlAn4 | -0.0413 | 0.0113 | 153 | -0.0681 | -0.0146 | -3.657 | 0.0010 |
| recall | | MetaPhlAn4 - Kraken2 | 0.1817 | 0.0102 | 153 | 0.1575 | 0.2059 | 17.761 | < 0.0001 |
| recall | | THMCv2 - Kraken2 | 0.6337 | 0.0102 | 153 | 0.6095 | 0.6580 | 61.946 | < 0.0001 |
| recall | | THMCv2 - MetaPhlAn4 | 0.4520 | 0.0102 | 153 | 0.4278 | 0.4763 | 44.185 | < 0.0001 |
| F1 | | MetaPhlAn4 - Kraken2 | 0.3233 | 0.0097 | 153 | 0.3003 | 0.3464 | 33.206 | < 0.0001 |
| F1 | | THMCv2 - Kraken2 | 0.5760 | 0.0097 | 153 | 0.5530 | 0.5991 | 59.153 | < 0.0001 |
| F1 | | THMCv2 - MetaPhlAn4 | 0.2527 | 0.0097 | 153 | 0.2296 | 0.2757 | 25.947 | < 0.0001 |
| AUPR | | MetaPhlAn4 - Kraken2 | 0.2411 | 0.0106 | 153 | 0.2159 | 0.2662 | 22.679 | < 0.0001 |
| AUPR | | THMCv2 - Kraken2 | 0.6922 | 0.0106 | 153 | 0.6670 | 0.7173 | 65.114 | < 0.0001 |
| AUPR | | THMCv2 - MetaPhlAn4 | 0.4511 | 0.0106 | 153 | 0.4259 | 0.4762 | 42.434 | < 0.0001 |
| precision | Stool Child | MetaPhlAn4 - Kraken2 | 0.5736 | 0.0130 | 147 | 0.5428 | 0.6043 | 44.170 | < 0.0001 |
| precision | | THMCv2 - Kraken2 | 0.5198 | 0.0130 | 147 | 0.4890 | 0.5505 | 40.027 | < 0.0001 |
| precision | | THMCv2 - MetaPhlAn4 | -0.0538 | 0.0130 | 147 | -0.0845 | -0.0230 | -4.143 | 0.0002 |
| recall | | MetaPhlAn4 - Kraken2 | 0.1970 | 0.0094 | 147 | 0.1749 | 0.2192 | 21.068 | < 0.0001 |
| recall | | THMCv2 - Kraken2 | 0.6327 | 0.0094 | 147 | 0.6106 | 0.6549 | 67.659 | < 0.0001 |
| recall | | THMCv2 - MetaPhlAn4 | 0.4357 | 0.0094 | 147 | 0.4136 | 0.4579 | 46.591 | < 0.0001 |
| F1 | | MetaPhlAn4 - Kraken2 | 0.3433 | 0.0098 | 147 | 0.3202 | 0.3664 | 35.178 | < 0.0001 |
| F1 | | THMCv2 - Kraken2 | 0.5777 | 0.0098 | 147 | 0.5546 | 0.6008 | 59.193 | < 0.0001 |
| F1 | | THMCv2 - MetaPhlAn4 | 0.2344 | 0.0098 | 147 | 0.2113 | 0.2575 | 24.015 | < 0.0001 |
| AUPR | | MetaPhlAn4 - Kraken2 | 0.2534 | 0.0102 | 147 | 0.2293 | 0.2774 | 24.946 | < 0.0001 |
| AUPR | | THMCv2 - Kraken2 | 0.6834 | 0.0102 | 147 | 0.6593 | 0.7074 | 67.283 | < 0.0001 |
| AUPR | | THMCv2 - MetaPhlAn4 | 0.4300 | 0.0102 | 147 | 0.4060 | 0.4541 | 42.337 | < 0.0001 |
| precision | Stool Toddler | MetaPhlAn4 - Kraken2 | 0.5777 | 0.0127 | 147 | 0.5477 | 0.6077 | 45.605 | < 0.0001 |
| precision | | THMCv2 - Kraken2 | 0.5038 | 0.0127 | 147 | 0.4738 | 0.5338 | 39.769 | < 0.0001 |
| precision | | THMCv2 - MetaPhlAn4 | -0.0739 | 0.0127 | 147 | -0.1039 | -0.0439 | -5.837 | 9.80e-08 |
| recall | | MetaPhlAn4 - Kraken2 | 0.1648 | 0.0118 | 147 | 0.1368 | 0.1929 | 13.918 | < 0.0001 |
| recall | | THMCv2 - Kraken2 | 0.6017 | 0.0118 | 147 | 0.5737 | 0.6298 | 50.813 | < 0.0001 |
| recall | | THMCv2 - MetaPhlAn4 | 0.4369 | 0.0118 | 147 | 0.4089 | 0.4650 | 36.895 | < 0.0001 |
| F1 | | MetaPhlAn4 - Kraken2 | 0.3256 | 0.0114 | 147 | 0.2985 | 0.3527 | 28.478 | < 0.0001 |
| F1 | | THMCv2 - Kraken2 | 0.5520 | 0.0114 | 147 | 0.5250 | 0.5791 | 48.282 | < 0.0001 |
| F1 | | THMCv2 - MetaPhlAn4 | 0.2264 | 0.0114 | 147 | 0.1993 | 0.2535 | 19.803 | < 0.0001 |
| AUPR | | MetaPhlAn4 - Kraken2 | 0.2267 | 0.0121 | 147 | 0.1981 | 0.2553 | 18.764 | < 0.0001 |
| AUPR | | THMCv2 - Kraken2 | 0.6524 | 0.0121 | 147 | 0.6238 | 0.6810 | 53.989 | < 0.0001 |
| AUPR | | THMCv2 - MetaPhlAn4 | 0.4256 | 0.0121 | 147 | 0.3970 | 0.4542 | 35.225 | < 0.0001 |
| precision | Stool Baby > 6 mo | MetaPhlAn4 - Kraken2 | 0.6131 | 0.0163 | 147 | 0.5746 | 0.6516 | 37.683 | < 0.0001 |

| Metric | Group | Contrast | Estimate | SE | df | 95% CI low | 95% CI high | t | adj. p |
|---|---|---|---|---|---|---|---|---|---|
| precision | | THMCv2 - Kraken2 | 0.4988 | 0.0163 | 147 | 0.4603 | 0.5373 | 30.657 | < 0.0001 |
| precision | | THMCv2 - MetaPhlAn4 | -0.1143 | 0.0163 | 147 | -0.1528 | -0.0758 | -7.026 | 2.21e-10 |
| recall | | MetaPhlAn4 - Kraken2 | 0.1100 | 0.0159 | 147 | 0.0724 | 0.1477 | 6.920 | 3.89e-10 |
| recall | | THMCv2 - Kraken2 | 0.5655 | 0.0159 | 147 | 0.5278 | 0.6031 | 35.565 | < 0.0001 |
| recall | | THMCv2 - MetaPhlAn4 | 0.4554 | 0.0159 | 147 | 0.4178 | 0.4931 | 28.645 | < 0.0001 |
| F1 | | MetaPhlAn4 - Kraken2 | 0.3220 | 0.0147 | 147 | 0.2872 | 0.3569 | 21.886 | < 0.0001 |
| F1 | | THMCv2 - Kraken2 | 0.5402 | 0.0147 | 147 | 0.5053 | 0.5750 | 36.710 | < 0.0001 |
| F1 | | THMCv2 - MetaPhlAn4 | 0.2181 | 0.0147 | 147 | 0.1833 | 0.2530 | 14.824 | < 0.0001 |
| AUPR | | MetaPhlAn4 - Kraken2 | 0.2003 | 0.0153 | 147 | 0.1641 | 0.2364 | 13.119 | < 0.0001 |
| AUPR | | THMCv2 - Kraken2 | 0.6178 | 0.0153 | 147 | 0.5816 | 0.6539 | 40.464 | < 0.0001 |
| AUPR | | THMCv2 - MetaPhlAn4 | 0.4175 | 0.0153 | 147 | 0.3813 | 0.4536 | 27.346 | < 0.0001 |
| precision | Stool Baby < 6 mo | MetaPhlAn4 - Kraken2 | 0.6242 | 0.0247 | 147 | 0.5657 | 0.6827 | 25.261 | < 0.0001 |
| precision | | THMCv2 - Kraken2 | 0.4405 | 0.0247 | 147 | 0.3820 | 0.4990 | 17.828 | < 0.0001 |
| precision | | THMCv2 - MetaPhlAn4 | -0.1837 | 0.0247 | 147 | -0.2422 | -0.1252 | -7.433 | < 0.0001 |
| recall | | MetaPhlAn4 - Kraken2 | 0.0562 | 0.0206 | 147 | 0.0073 | 0.1050 | 2.721 | 0.0198 |
| recall | | THMCv2 - Kraken2 | 0.5004 | 0.0206 | 147 | 0.4516 | 0.5493 | 24.250 | < 0.0001 |
| recall | | THMCv2 - MetaPhlAn4 | 0.4443 | 0.0206 | 147 | 0.3954 | 0.4931 | 21.529 | < 0.0001 |
| F1 | | MetaPhlAn4 - Kraken2 | 0.3280 | 0.0224 | 147 | 0.2750 | 0.3810 | 14.646 | < 0.0001 |
| F1 | | THMCv2 - Kraken2 | 0.4898 | 0.0224 | 147 | 0.4368 | 0.5428 | 21.870 | < 0.0001 |
| F1 | | THMCv2 - MetaPhlAn4 | 0.1618 | 0.0224 | 147 | 0.1088 | 0.2148 | 7.224 | < 0.0001 |
| AUPR | | MetaPhlAn4 - Kraken2 | 0.1608 | 0.0216 | 147 | 0.1095 | 0.2121 | 7.428 | < 0.0001 |
| AUPR | | THMCv2 - Kraken2 | 0.5216 | 0.0216 | 147 | 0.4703 | 0.5728 | 24.095 | < 0.0001 |
| AUPR | | THMCv2 - MetaPhlAn4 | 0.3608 | 0.0216 | 147 | 0.3095 | 0.4120 | 16.667 | < 0.0001 |
| precision | Vaginal Adult | MetaPhlAn4 - Kraken2 | 0.5902 | 0.0230 | 141 | 0.5357 | 0.6448 | 25.628 | < 0.0001 |
| precision | | THMCv2 - Kraken2 | 0.5226 | 0.0230 | 141 | 0.4680 | 0.5771 | 22.690 | < 0.0001 |
| precision | | THMCv2 - MetaPhlAn4 | -0.0677 | 0.0230 | 141 | -0.1222 | -0.0131 | -2.938 | 0.0107 |
| recall | | MetaPhlAn4 - Kraken2 | 0.3399 | 0.0254 | 141 | 0.2798 | 0.4000 | 13.397 | < 0.0001 |
| recall | | THMCv2 - Kraken2 | 0.5609 | 0.0254 | 141 | 0.5008 | 0.6210 | 22.109 | < 0.0001 |
| recall | | THMCv2 - MetaPhlAn4 | 0.2210 | 0.0254 | 141 | 0.1609 | 0.2811 | 8.712 | < 0.0001 |
| F1 | | MetaPhlAn4 - Kraken2 | 0.4673 | 0.0205 | 141 | 0.4186 | 0.5159 | 22.741 | < 0.0001 |
| F1 | | THMCv2 - Kraken2 | 0.5584 | 0.0205 | 141 | 0.5097 | 0.6071 | 27.176 | < 0.0001 |
| F1 | | THMCv2 - MetaPhlAn4 | 0.0911 | 0.0205 | 141 | 0.0424 | 0.1398 | 4.435 | 5.45e-05 |
| AUPR | | MetaPhlAn4 - Kraken2 | 0.4256 | 0.0241 | 141 | 0.3686 | 0.4826 | 17.696 | < 0.0001 |
| AUPR | | THMCv2 - Kraken2 | 0.6408 | 0.0241 | 141 | 0.5838 | 0.6978 | 26.644 | < 0.0001 |
| AUPR | | THMCv2 - MetaPhlAn4 | 0.2152 | 0.0241 | 141 | 0.1582 | 0.2722 | 8.948 | < 0.0001 |

**Table S2.** Pairwise comparisons of abundance-weighted classification performance metrics across methods, grouped by body site and age group. Estimates reflect mean differences between methods. Adjusted p-values were corrected for multiple comparisons. MSE, mean squared error; SE, standard error; CI, confidence interval; df, degrees of freedom.

| Metric | Group | Contrast | Estimate | SE | df | 95% CI low | 95% CI high | t | adj. p |
|---|---|---|---|---|---|---|---|---|---|
| precision | Stool Adult | MetaPhlAn4 - Kraken2 | 0.109 | 0.015 | 153 | 0.073 | 0.144 | 7.228 | < 0.0001 |
| precision | | THMCv2 - Kraken2 | 0.156 | 0.015 | 153 | 0.121 | 0.192 | 10.385 | < 0.0001 |
| precision | | THMCv2 - MetaPhlAn4 | 0.047 | 0.015 | 153 | 0.012 | 0.083 | 3.157 | 0.0050 |
| recall | | MetaPhlAn4 - Kraken2 | 0.329 | 0.017 | 153 | 0.289 | 0.368 | 19.698 | < 0.0001 |
| recall | | THMCv2 - Kraken2 | 0.341 | 0.017 | 153 | 0.302 | 0.381 | 20.434 | < 0.0001 |
| recall | | THMCv2 - MetaPhlAn4 | 0.012 | 0.017 | 153 | -0.027 | 0.052 | 0.736 | 0.7430 |
| F1 | | MetaPhlAn4 - Kraken2 | 0.231 | 0.015 | 153 | 0.195 | 0.268 | 15.157 | < 0.0001 |
| F1 | | THMCv2 - Kraken2 | 0.263 | 0.015 | 153 | 0.227 | 0.299 | 17.215 | < 0.0001 |
| F1 | | THMCv2 - MetaPhlAn4 | 0.031 | 0.015 | 153 | -0.005 | 0.068 | 2.058 | 0.1020 |
| AUPR | | MetaPhlAn4 - Kraken2 | 0.334 | 0.026 | 153 | 0.272 | 0.396 | 12.698 | < 0.0001 |
| AUPR | | THMCv2 - Kraken2 | 0.393 | 0.026 | 153 | 0.330 | 0.455 | 14.936 | < 0.0001 |
| AUPR | | THMCv2 - MetaPhlAn4 | 0.059 | 0.026 | 153 | -0.003 | 0.121 | 2.238 | 0.0680 |
| precision | Stool Child | MetaPhlAn4 - Kraken2 | 0.075 | 0.013 | 147 | 0.045 | 0.105 | 5.947 | < 0.0001 |
| precision | | THMCv2 - Kraken2 | 0.119 | 0.013 | 147 | 0.089 | 0.149 | 9.390 | < 0.0001 |
| precision | | THMCv2 - MetaPhlAn4 | 0.044 | 0.013 | 147 | 0.014 | 0.073 | 3.444 | 0.0020 |
| recall | | MetaPhlAn4 - Kraken2 | 0.310 | 0.016 | 147 | 0.272 | 0.348 | 19.280 | < 0.0001 |
| recall | | THMCv2 - Kraken2 | 0.321 | 0.016 | 147 | 0.283 | 0.359 | 19.968 | < 0.0001 |
| recall | | THMCv2 - MetaPhlAn4 | 0.011 | 0.016 | 147 | -0.027 | 0.049 | 0.688 | 0.7710 |
| F1 | | MetaPhlAn4 - Kraken2 | 0.208 | 0.013 | 147 | 0.176 | 0.240 | 15.443 | < 0.0001 |
| F1 | | THMCv2 - Kraken2 | 0.237 | 0.013 | 147 | 0.205 | 0.269 | 17.550 | < 0.0001 |
| F1 | | THMCv2 - MetaPhlAn4 | 0.028 | 0.013 | 147 | -0.004 | 0.060 | 2.107 | 0.0920 |
| AUPR | | MetaPhlAn4 - Kraken2 | 0.296 | 0.022 | 147 | 0.243 | 0.350 | 13.232 | < 0.0001 |
| AUPR | | THMCv2 - Kraken2 | 0.347 | 0.022 | 147 | 0.294 | 0.400 | 15.501 | < 0.0001 |
| AUPR | | THMCv2 - MetaPhlAn4 | 0.051 | 0.022 | 147 | -0.002 | 0.104 | 2.269 | 0.0630 |
| precision | Stool Toddler | MetaPhlAn4 - Kraken2 | 0.084 | 0.016 | 147 | 0.045 | 0.123 | 5.152 | < 0.0001 |
| precision | | THMCv2 - Kraken2 | 0.124 | 0.016 | 147 | 0.085 | 0.162 | 7.577 | < 0.0001 |
| precision | | THMCv2 - MetaPhlAn4 | 0.040 | 0.016 | 147 | 0.001 | 0.078 | 2.426 | 0.0430 |
| recall | | MetaPhlAn4 - Kraken2 | 0.269 | 0.017 | 147 | 0.228 | 0.309 | 15.692 | < 0.0001 |
| recall | | THMCv2 - Kraken2 | 0.279 | 0.017 | 147 | 0.239 | 0.320 | 16.301 | < 0.0001 |
| recall | | THMCv2 - MetaPhlAn4 | 0.010 | 0.017 | 147 | -0.030 | 0.051 | 0.609 | 0.8160 |
| F1 | | MetaPhlAn4 - Kraken2 | 0.186 | 0.016 | 147 | 0.149 | 0.223 | 11.992 | < 0.0001 |
| F1 | | THMCv2 - Kraken2 | 0.212 | 0.016 | 147 | 0.176 | 0.249 | 13.686 | < 0.0001 |
| F1 | | THMCv2 - MetaPhlAn4 | 0.026 | 0.016 | 147 | -0.010 | 0.063 | 1.694 | 0.2110 |
| AUPR | | MetaPhlAn4 - Kraken2 | 0.281 | 0.024 | 147 | 0.223 | 0.338 | 11.526 | < 0.0001 |
| AUPR | | THMCv2 - Kraken2 | 0.317 | 0.024 | 147 | 0.260 | 0.375 | 13.038 | < 0.0001 |
| AUPR | | THMCv2 - MetaPhlAn4 | 0.037 | 0.024 | 147 | -0.021 | 0.094 | 1.512 | 0.2880 |
| precision | Stool Baby > 6 mo | MetaPhlAn4 - Kraken2 | 0.054 | 0.013 | 147 | 0.025 | 0.084 | 4.321 | < 0.0001 |

| Metric | Group | Contrast | Estimate | SE | df | 95% CI low | 95% CI high | t | adj. p |
|---|---|---|---|---|---|---|---|---|---|
| precision | | THMCv2 - Kraken2 | 0.087 | 0.013 | 147 | 0.057 | 0.117 | 6.893 | < 0.0001 |
| precision | | THMCv2 - MetaPhlAn4 | 0.032 | 0.013 | 147 | 0.003 | 0.062 | 2.572 | 0.0300 |
| recall | | MetaPhlAn4 - Kraken2 | 0.236 | 0.017 | 147 | 0.196 | 0.276 | 14.019 | < 0.0001 |
| recall | | THMCv2 - Kraken2 | 0.242 | 0.017 | 147 | 0.202 | 0.282 | 14.363 | < 0.0001 |
| recall | | THMCv2 - MetaPhlAn4 | 0.006 | 0.017 | 147 | -0.034 | 0.046 | 0.344 | 0.9370 |
| F1 | | MetaPhlAn4 - Kraken2 | 0.156 | 0.013 | 147 | 0.124 | 0.188 | 11.552 | < 0.0001 |
| F1 | | THMCv2 - Kraken2 | 0.176 | 0.013 | 147 | 0.144 | 0.208 | 13.063 | < 0.0001 |
| F1 | | THMCv2 - MetaPhlAn4 | 0.020 | 0.013 | 147 | -0.012 | 0.052 | 1.512 | 0.2880 |
| AUPR | | MetaPhlAn4 - Kraken2 | 0.220 | 0.021 | 147 | 0.170 | 0.270 | 10.447 | < 0.0001 |
| AUPR | | THMCv2 - Kraken2 | 0.254 | 0.021 | 147 | 0.204 | 0.304 | 12.074 | < 0.0001 |
| AUPR | | THMCv2 - MetaPhlAn4 | 0.034 | 0.021 | 147 | -0.016 | 0.084 | 1.627 | 0.2380 |
| precision | Stool Baby < 6 mo | MetaPhlAn4 - Kraken2 | 0.084 | 0.022 | 147 | 0.032 | 0.136 | 3.815 | 0.0010 |
| precision | | THMCv2 - Kraken2 | 0.105 | 0.022 | 147 | 0.053 | 0.158 | 4.779 | < 0.0001 |
| precision | | THMCv2 - MetaPhlAn4 | 0.021 | 0.022 | 147 | -0.031 | 0.073 | 0.964 | 0.6010 |
| recall | | MetaPhlAn4 - Kraken2 | 0.188 | 0.024 | 147 | 0.130 | 0.246 | 7.701 | < 0.0001 |
| recall | | THMCv2 - Kraken2 | 0.201 | 0.024 | 147 | 0.143 | 0.258 | 8.217 | < 0.0001 |
| recall | | THMCv2 - MetaPhlAn4 | 0.013 | 0.024 | 147 | -0.045 | 0.070 | 0.516 | 0.8640 |
| F1 | | MetaPhlAn4 - Kraken2 | 0.140 | 0.022 | 147 | 0.087 | 0.193 | 6.271 | < 0.0001 |
| F1 | | THMCv2 - Kraken2 | 0.161 | 0.022 | 147 | 0.108 | 0.214 | 7.206 | < 0.0001 |
| F1 | | THMCv2 - MetaPhlAn4 | 0.021 | 0.022 | 147 | -0.032 | 0.074 | 0.934 | 0.6190 |
| AUPR | | MetaPhlAn4 - Kraken2 | 0.188 | 0.032 | 147 | 0.113 | 0.263 | 5.935 | < 0.0001 |
| AUPR | | THMCv2 - Kraken2 | 0.225 | 0.032 | 147 | 0.150 | 0.300 | 7.111 | < 0.0001 |
| AUPR | | THMCv2 - MetaPhlAn4 | 0.037 | 0.032 | 147 | -0.038 | 0.112 | 1.177 | 0.4690 |
| precision | Vaginal Adult | MetaPhlAn4 - Kraken2 | 0.117 | 0.018 | 141 | 0.074 | 0.160 | 6.463 | < 0.0001 |
| precision | | THMCv2 - Kraken2 | 0.117 | 0.018 | 141 | 0.075 | 0.160 | 6.486 | < 0.0001 |
| precision | | THMCv2 - MetaPhlAn4 | 0.000 | 0.018 | 141 | -0.042 | 0.043 | 0.023 | 1.0000 |
| recall | | MetaPhlAn4 - Kraken2 | 0.211 | 0.023 | 141 | 0.157 | 0.264 | 9.271 | < 0.0001 |
| recall | | THMCv2 - Kraken2 | 0.213 | 0.023 | 141 | 0.159 | 0.267 | 9.373 | < 0.0001 |
| recall | | THMCv2 - MetaPhlAn4 | 0.002 | 0.023 | 141 | -0.051 | 0.056 | 0.101 | 0.9940 |
| F1 | | MetaPhlAn4 - Kraken2 | 0.178 | 0.020 | 141 | 0.132 | 0.225 | 9.047 | < 0.0001 |
| F1 | | THMCv2 - Kraken2 | 0.180 | 0.020 | 141 | 0.133 | 0.226 | 9.116 | < 0.0001 |
| F1 | | THMCv2 - MetaPhlAn4 | 0.001 | 0.020 | 141 | -0.045 | 0.048 | 0.069 | 0.9970 |
| AUPR | | MetaPhlAn4 - Kraken2 | 0.252 | 0.029 | 141 | 0.183 | 0.322 | 8.584 | < 0.0001 |
| AUPR | | THMCv2 - Kraken2 | 0.255 | 0.029 | 141 | 0.185 | 0.324 | 8.665 | < 0.0001 |
| AUPR | | THMCv2 - MetaPhlAn4 | 0.002 | 0.029 | 141 | -0.067 | 0.072 | 0.082 | 0.9960 |

**Table S3.** Pairwise comparisons of mean taxon abundance among false negatives (FN) and false positives (FP) across methods, grouped by body site and age group. Estimates reflect mean differences between methods. Adjusted p-values were corrected for multiple comparisons. Abund. FN, mean abundance of false negatives; Abund. FP, mean abundance of false positives; SE, standard error; CI, confidence interval; df, degrees of freedom.

| Metric | Group | Contrast | Estimate | SE | df | 95% CI low | 95% CI high | t | adj. p |
|---|---|---|---|---|---|---|---|---|---|
| Abund. FN | Stool Adult | MetaPhlAn4 - Kraken2 | -0.003 | 0.000 | 143 | -0.004 | -0.003 | -17.219 | < 0.0001 |
| Abund. FN | | THMCv2 - Kraken2 | -0.003 | 0.000 | 143 | -0.004 | -0.003 | -16.357 | < 0.0001 |
| Abund. FN | | THMCv2 - MetaPhlAn4 | 0.000 | 0.000 | 143 | 0.000 | 0.000 | -0.079 | 0.9970 |
| Abund. FP | | MetaPhlAn4 - Kraken2 | 0.004 | 0.001 | 153 | 0.002 | 0.006 | 5.878 | < 0.0001 |
| Abund. FP | | THMCv2 - Kraken2 | -0.001 | 0.001 | 153 | -0.003 | 0.001 | -1.462 | 0.3120 |
| Abund. FP | | THMCv2 - MetaPhlAn4 | -0.005 | 0.001 | 153 | -0.007 | -0.003 | -7.339 | < 0.0001 |
| Abund. FN | Stool Child | MetaPhlAn4 - Kraken2 | -0.003 | 0.000 | 139 | -0.004 | -0.003 | -12.076 | < 0.0001 |
| Abund. FN | | THMCv2 - Kraken2 | -0.003 | 0.000 | 139 | -0.004 | -0.003 | -11.667 | < 0.0001 |
| Abund. FN | | THMCv2 - MetaPhlAn4 | 0.000 | 0.000 | 139 | -0.001 | 0.001 | -0.128 | 0.9910 |
| Abund. FP | | MetaPhlAn4 - Kraken2 | 0.004 | 0.001 | 146 | 0.003 | 0.006 | 7.877 | < 0.0001 |
| Abund. FP | | THMCv2 - Kraken2 | -0.001 | 0.001 | 146 | -0.002 | 0.001 | -1.397 | 0.3450 |
| Abund. FP | | THMCv2 - MetaPhlAn4 | -0.005 | 0.001 | 146 | -0.006 | -0.004 | -9.268 | < 0.0001 |
| Abund. FN | Stool Toddler | MetaPhlAn4 - Kraken2 | -0.003 | 0.000 | 137 | -0.004 | -0.003 | -15.647 | < 0.0001 |
| Abund. FN | | THMCv2 - Kraken2 | -0.003 | 0.000 | 137 | -0.004 | -0.003 | -15.134 | < 0.0001 |
| Abund. FN | | THMCv2 - MetaPhlAn4 | 0.000 | 0.000 | 137 | -0.001 | 0.000 | -0.382 | 0.9230 |
| Abund. FP | | MetaPhlAn4 - Kraken2 | 0.008 | 0.002 | 146 | 0.004 | 0.011 | 5.052 | < 0.0001 |
| Abund. FP | | THMCv2 - Kraken2 | -0.001 | 0.002 | 146 | -0.005 | 0.003 | -0.655 | 0.7900 |
| Abund. FP | | THMCv2 - MetaPhlAn4 | -0.009 | 0.002 | 146 | -0.012 | -0.005 | -5.704 | < 0.0001 |
| Abund. FN | Stool Baby > 6 mo | MetaPhlAn4 - Kraken2 | -0.005 | 0.001 | 135 | -0.007 | -0.003 | -6.402 | < 0.0001 |
| Abund. FN | | THMCv2 - Kraken2 | -0.005 | 0.001 | 135 | -0.008 | -0.003 | -6.015 | < 0.0001 |
| Abund. FN | | THMCv2 - MetaPhlAn4 | 0.000 | 0.001 | 135 | -0.002 | 0.002 | -0.066 | 0.9980 |
| Abund. FP | | MetaPhlAn4 - Kraken2 | 0.011 | 0.003 | 145 | 0.005 | 0.017 | 4.399 | < 0.0001 |
| Abund. FP | | THMCv2 - Kraken2 | -0.001 | 0.003 | 145 | -0.007 | 0.005 | -0.225 | 0.9730 |
| Abund. FP | | THMCv2 - MetaPhlAn4 | -0.012 | 0.003 | 145 | -0.018 | -0.006 | -4.621 | < 0.0001 |
| Abund. FN | Stool Baby < 6 mo | MetaPhlAn4 - Kraken2 | -0.005 | 0.001 | 132 | -0.007 | -0.002 | -4.534 | < 0.0001 |
| Abund. FN | | THMCv2 - Kraken2 | -0.005 | 0.001 | 132 | -0.008 | -0.003 | -4.807 | < 0.0001 |
| Abund. FN | | THMCv2 - MetaPhlAn4 | -0.001 | 0.001 | 132 | -0.003 | 0.002 | -0.692 | 0.7690 |
| Abund. FP | | MetaPhlAn4 - Kraken2 | 0.018 | 0.011 | 141 | -0.008 | 0.044 | 1.613 | 0.2440 |
| Abund. FP | | THMCv2 - Kraken2 | -0.001 | 0.011 | 141 | -0.026 | 0.024 | -0.063 | 0.9980 |
| Abund. FP | | THMCv2 - MetaPhlAn4 | -0.018 | 0.011 | 141 | -0.044 | 0.008 | -1.674 | 0.2190 |
| Abund. FN | Vaginal Adult | MetaPhlAn4 - Kraken2 | -0.009 | 0.003 | 91 | -0.017 | -0.001 | -2.796 | 0.0170 |
| Abund. FN | | THMCv2 - Kraken2 | -0.010 | 0.005 | 91 | -0.023 | 0.003 | -1.748 | 0.1930 |
| Abund. FN | | THMCv2 - MetaPhlAn4 | 0.000 | 0.006 | 91 | -0.014 | 0.013 | -0.063 | 0.9980 |
| Abund. FP | | MetaPhlAn4 - Kraken2 | -0.001 | 0.000 | 128 | -0.001 | 0.000 | -2.053 | 0.1040 |
| Abund. FP | | THMCv2 - Kraken2 | -0.001 | 0.000 | 128 | -0.002 | -0.001 | -4.910 | < 0.0001 |
| Abund. FP | | THMCv2 - MetaPhlAn4 | -0.001 | 0.000 | 128 | -0.002 | 0.000 | -2.693 | 0.0220 |
